# How subunit rotation controls tRNA dynamics in the ribosome

**DOI:** 10.1101/2025.01.22.634288

**Authors:** Sandra Byju, Paul C. Whitford

## Abstract

In order to translate messenger RNA into proteins, the ribosome must coordinate a wide range of conformational rearrangements. Some steps involve individual molecules, whereas others require synchronization of multiple collective motions. For example, the ribosomal “small” subunit (∼1 MDa) is known to undergo rotational motion (∼ 10*^◦^*) that is correlated with large-scale displacements of tRNA molecules (∼50Å). While decades of biochemical, single-molecule and structural analysis have provided many insights into the timing of these motions, little is known about how these dynamical processes influence each other. To address this, we use molecular simulations to isolate specific interactions that allow tRNA kinetics to be controlled by subunit rotation. Specifically, we applied an all-atom structure-based model to perform molecular simulations of P/E hybrid formation, a process by which the tRNA molecules are displaced between ribosomal binding sites. By varying the extent of subunit rotation in each simulation, these calculations reveal a pronounced non-monotonic dependence of tRNA kinetics on subunit rotation. That is, tRNA kinetics increase and then decrease as the subunit is rotated. We further show that this behavior is due to the steric contributions of a single ribosomal protein. Together, these calculations provide insights into the physicochemical properties of the ribosome, while establishing a strategy for isolating causal relationships in biomolecular assemblies.

## Introduction

The ribosome is a massive nucleoprotein assembly that is responsible for protein synthesis in all living cells. In order to function, it coordinates the motion of multiple ribosomal RNA (rRNA) molecules and more than 50 proteins. Overall, the structure is generally described in terms of two subunits, called the small subunit (SSU) and large subunit (LSU; Fig. 1). In bacteria, the LSU contains the 23S and 5S rRNA molecules along with proteins, while the SSU is composed of the 16S rRNA and proteins. The ribosome, along with the transfer RNA (tRNA) molecules, performs the function of decoding the genetic information in messenger RNA (mRNA), in order to synthesize proteins (i.e., elongation). During elongation, each tRNA molecule sequentially binds the ribosomal A (aminoacyl), P (peptidyl), and E (exit) sites, where precise kinetic properties ensure accurate and efficient gene expression.

**Figure 1:**
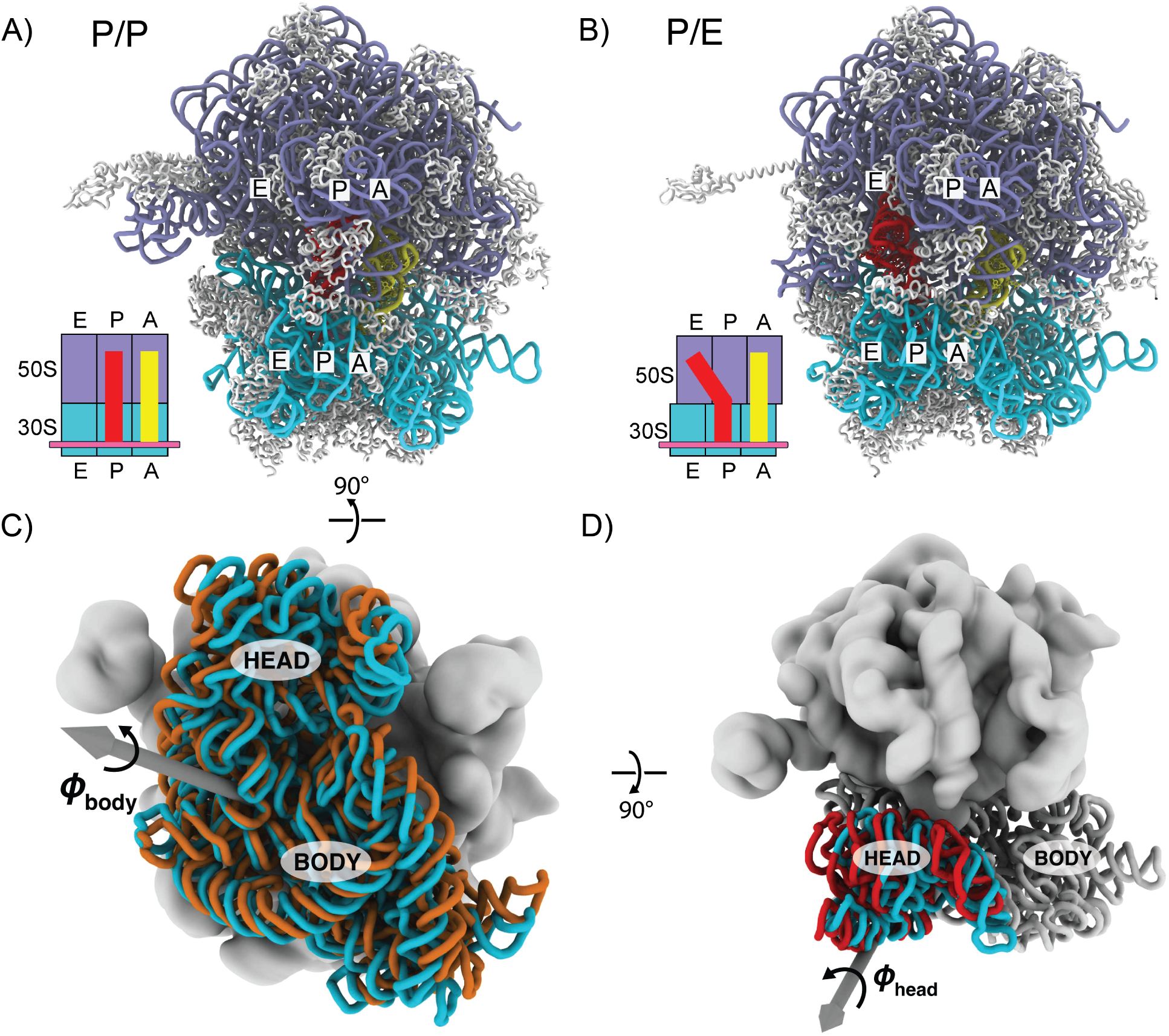
P/E tRNA hybrid formation and ribosomal subunit rotation. A) E. coli ribosome (PDB ID: 4V9D^43^) in the PRE-translocation state, with ribosome in the unrotated configuration. The A-site (yellow) and P-site tRNAs (red) are in the A/A and P/P states, respectively. B) E. coli ribosome in the P/E tRNA hybrid state, where the ribosome is in a rotated configuration.^43^ The LSU rRNA is shown in violet, SSU rRNA in cyan and proteins in white. C-D) The SSU body and head domains undergo multiple modes of rotation during elongation. We use the RAD method^41^ to decompose the body and head domain orientations. *φ*_body_ describes rotation of the SSU body, relative to LSU (gray; panel C). *φ*_head_ describes rotation of the SSU head, relative to the SSU body (panel D).

During the elongation cycle, the ribosome and tRNA molecules undergo several large-scale conformational rearrangements.^1–4^ After the decoding step, where elongation factor Tu (EF-Tu) delivers the aminoacyl-tRNA (aa-tRNA) to the A site, the ribosome facilitates accommodation of aa-tRNA into the peptidyl transferase center (PTC). At the PTC, a peptide bond is formed when the nascent peptide chain is transferred from the peptidyl tRNA to the aa-tRNA, resulting in the pre-translocation state (A/A-P/P). This is followed by the translocation step, where tRNAs in the A and P sites transition to the P and E sites, respectively, and the mRNA advances by one codon. This full process occurs in two stages: P/E hybrid state formation on the large subunit (LSU) and mRNA-tRNA translocation on the small subunit (SSU). During hybrid state formation of the tRNA,^5^ there is concomitant rotation of the SSU body.^6^ Translocation is associated with the movement of mRNA and tRNA on the SSU, which is impacted by SSU head rotation, ^7^ tilting,^8^ and elongation factor G (EF-G). In the post-translocation state (P/P-E/E) the A site is vacant and ready for the next cycle of elongation to occur.

While there have been many efforts to explore the relationship between SSU body rotation and the kinetics of P/E hybrid formation, our physical-chemical understanding remains less developed. P/E tRNA hybrid formation involves the transition of deacylated P-site tRNA to the P/E hybrid state (Fig. 1B), and it is closely coupled to inter-subunit rotation^6,9^ (often referred to as “ratcheting”). By introducing a disulfide linker between the SSU and LSU, Horan et al.,^10^ provided direct evidence that inter-subunit rotation is required for translocation, while Ermolenko et al.,^11^ used FRET measurements to show that P/E hybrid formation is coupled to inter-subunit rotation in the ribosome. Consistent with these experiments, simulations also found that rotation was required for P/E formation to occur. ^12^ Upon rotation, the tRNA is able to interact with the L1 stalk, which further stabilizes the P/E state.^13–15^ While these studies forcefully demonstrated that hybrid formation is linked to subunit rotation, the precise interplay between SSU body rotation and the kinetics of P/E hybrid formation has not been characterized.

In this study, we use molecular dynamics simulations with a multi-basin all-atom structure-based model^8,16^ to investigate how SSU body rotation influences the kinetics of P/E tRNA hybrid formation. By simulating hybrid formation for different SSU rotation states, we identify the molecular factors that lead to rotation-dependent tRNA kinetics. This reveals a non-monotonic dependence of hybrid formation kinetics on SSU rotation, which originates from the changing steric character of the ribosome. Together, these calculations provide precise insights into the dynamics of P/E formation, while serving as an example for how to isolate the functional dependence between collective processes in biomolecular assemblies.

## Results

### Simulating hybrid formation for different SSU body rotation states

To investigate how the kinetics of P/E tRNA hybrid formation depends on small subunit body rotation, we used an all-atom structure-based (SMOG) model ^16^ to perform molecular dynamics simulations of the ribosome. Specifically, we simulated P/E hybrid formation events with the SSU at varied degrees of rotation. Since the model has all-atom resolution and simplified energetics, one can isolate how changes in the steric character of the ribosome control tRNA dynamics.^17^ In a conventional structure-based model, an experimental structure is explicitly defined to be the global potential energy minimum. ^18–20^ All other interactions are repulsive, which ensures that the steric composition is properly represented. These single-basin models have been particularly effective for studying biomolecular folding and conformational dynamics near a single native state.^21–23^ In addition, similar models with native interactions encoded have provided insights into ribosome assembly^24^ and nascent protein folding on the ribosome.^25–27^ When studying conformational transitions between multiple stable conformations, multi-basin approaches are more suitable. ^12,28^ Accordingly, in the current study we designed and applied a multi-basin SMOG model, where the pre-translocation (Fig. 1A) and post-translocation (Fig. S1) tRNA positions are defined as potential energy minima (see methods). With this representation, we simulated spontaneous (i.e., not targeted) transitions of the tRNA between the P/P and P/E conformations (Fig. 1). To address the relationship between rotation and tRNA movement, we repeated the simulations using different energetic representations of the SSU, such that specific orientations (rotation angles) were favored. Since only the endpoints are defined to be stable, any free-energy barriers that are observed may be attributed to changes in conformational entropy and steric interactions. As described below, we find that these contributions lead to an intricate relationship between rotation and tRNA dynamics.

To define our multi-basin SMOG models, we first assessed all experimentally determined structures of the E. coli ribosome (resolution of 8Å or better; 412 structures; Fig. 2A). From this, we identified representative conformations at different degrees of body rotation (*φ*_body_). To isolate the impact of body rotation on tRNA dynamics, we only considered structures in which head rotation was small (Fig. 2A; see methods for full details). From this, we selected seven structures for which *φ*_body_ ranges from 0*^◦^* to ∼ 11*^◦^*. We next used the representative experimental structures to construct SMOG models in which specific orientations of the SSU are favored. That is, tRNA-ribosome interactions are identical in all models, while the stabilizing intra-ribosome interactions in each model were defined based on a different representative structure (see Methods for details on force field construction). When performing simulations with these models, the mean body rotation angle, (*φ*_body_), was consistent with the associated representative structure (Fig. 2B, Table S1). For ease of notation, in subsequent discussion we will refer to simulations with each rotation model by the nearest integer value of the angle (0*^◦^*, 3*^◦^*, 4*^◦^*, 6*^◦^*, 7*^◦^*, 9*^◦^*, 11*^◦^*; Table S1).

**Figure 2:**
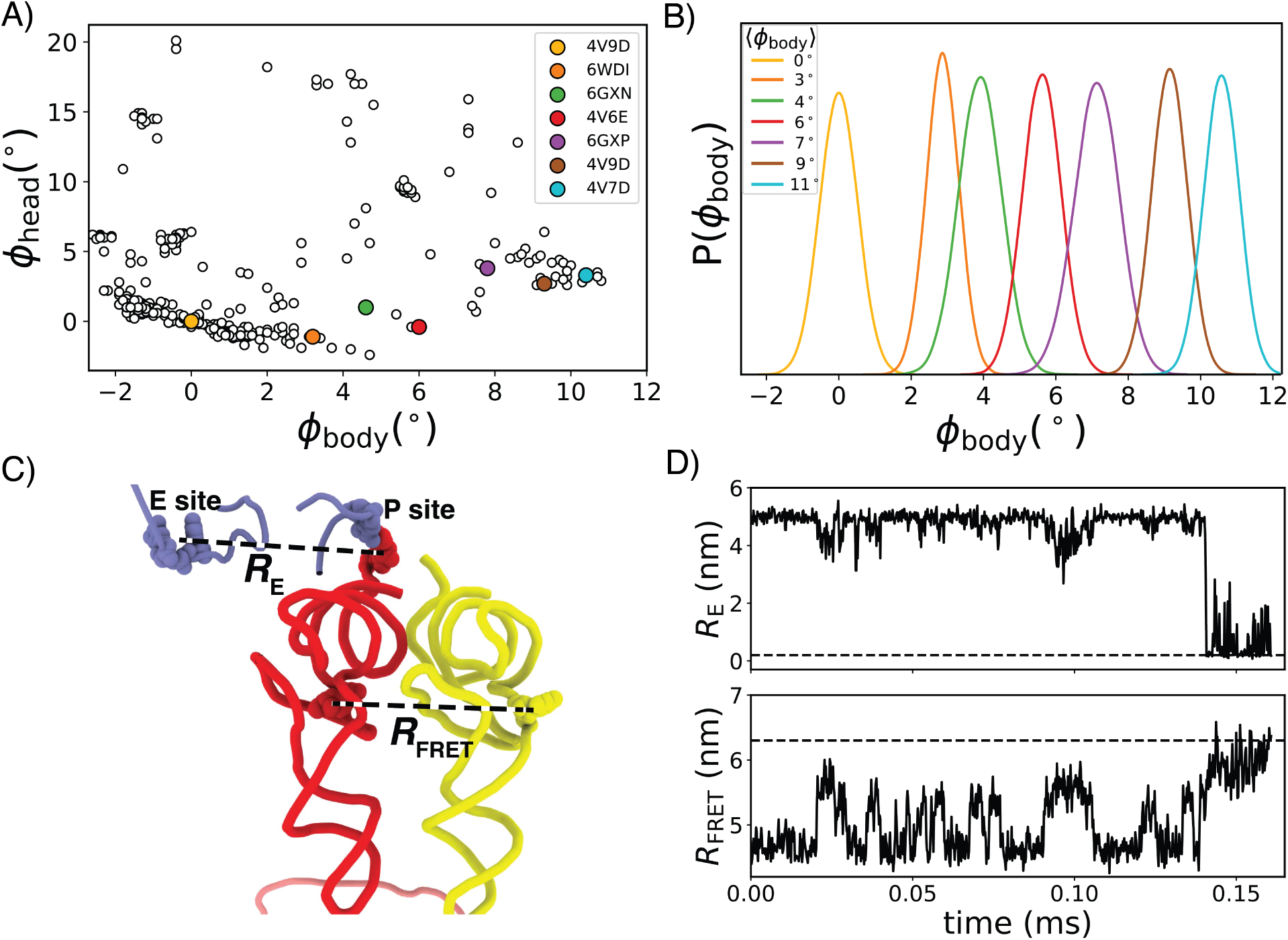
Simulating P/E tRNA hybrid formation for different body rotation angles. A) All E. coli ribosome structures in the RCSB database (resolution of 8Å or greater; N=412) is shown as a function of body rotation (*φ*_body_) and head rotation (*φ*_head_). Representative structures at varying *φ*_body_ (and small *φ*_head_) were used to generate the rotation force fields (see methods for details on selection criteria). The representative rotation structures are marked by colored circles. PDB IDs given in the legend. B) The probability distribution of *φ*_body_, calculated from simulations with each rotation model. The mean body rotation angle (*φ*_body_) is given in the legend. Colors are the same as panel A. C) To monitor the position of the P-site tRNA during hybrid formation, the distance between the 3’-CCA tail of the tRNA and the E site on the LSU (*R*_E_), as well as the distance between the elbows (*R*_FRET_), were calculated (see Methods). Structure colored as in Figs. 1A. D) Representative time trace for simulation with the 6*^◦^* rotation model. The P/E state is denoted by dashed lines as corresponds to *R*_E_ < 0.2nm and *R*_FRET_ > 6.3nm. Adoption of the P/E state is associated with large changes in both distances.

To quantify the relationship between P/E formation kinetics and body rotation, we performed many independent simulations with each model (707, in total). To determine when the tRNA reached the P/E state, we calculated the distance between the P-site tRNA 3’-CCA tail and the E site on the LSU (*R*_E_), as well as the distance between the elbow regions of P-site tRNA and A-site tRNA (*R*_FRET_; Figs. 2C and 2D; see methods for details). The label *R*_FRET_ is used since it is the distance between residues that are commonly probed in single-molecule FRET experiments.^29,30^ Consistent with biochemical measurements,^10^ P/E formation did not occur in any simulation where the ribosome was unrotated (Table S2). For intermediate values of the rotation angle (6*^◦^* and 7*^◦^*) every simulation reached the P/E state. Surprisingly, we find there is a slow down when the SSU is highly rotated (*>* 7*^◦^*), where hybrid state formation did not occur in many of the simulations. As described below, this indicates that rotation is associated with competing factors that pull the tRNA and impede its motion.

### Intermediate rotation exhibits fastest tRNA kinetics

While experimental studies have indicated that subunit rotation facilitates hybrid state formation,^11,13,14^ we find that the time required to reach the P/E state (*τ*) depends nonmonotonically on SSU rotation. As noted above, when the SSU was unrotated (*φ*_body_ = 0*^◦^*), P/E hybrid formation was not observed within the longest simulated timescale (8 ms). For *φ*_body_ = 3*^◦^*, transitions to the P/E state were observed in roughly half of the simulated trajectories (32 of 70; Fig. 3A), where each simulation was continued for up to 4 ms. From this, we obtained a lower bound of 2.9 ms for the mean first passage time *τ̄*. To calculate this lower bound, a time of 4 ms was used for each simulation that did not reach the hybrid state. When the body was further rotated to 4*^◦^*, the P/E conformation was reached in almost every simulation (118 of 119; Fig S2A). In our model, hybrid state formation is fastest when *φ*_body_ = 6*^◦^* (Fig. 3B). For this angle, all runs reached the P/E state, *τ̄* ∼ 0.2 ms, and the majority of simulated transitions (154 of 160) occurred in less than 0.5 ms. Similar kinetics were observed for *φ*_body_ = 7*^◦^* (Table S2; Fig S2B). For body rotation angles greater than 7*^◦^*, we again found that many of the simulations did not reach the P/E ensemble within the simulated time. Accordingly, there is a striking increase in *τ̄* for larger values of *φ*_body_. For example, at 11*^◦^* rotation, *τ̄* ∼ 2.3 ms, which represents at least an eleven-fold reduction of the kinetics, relative to the kinetics at 6*^◦^*.

**Figure 3:**
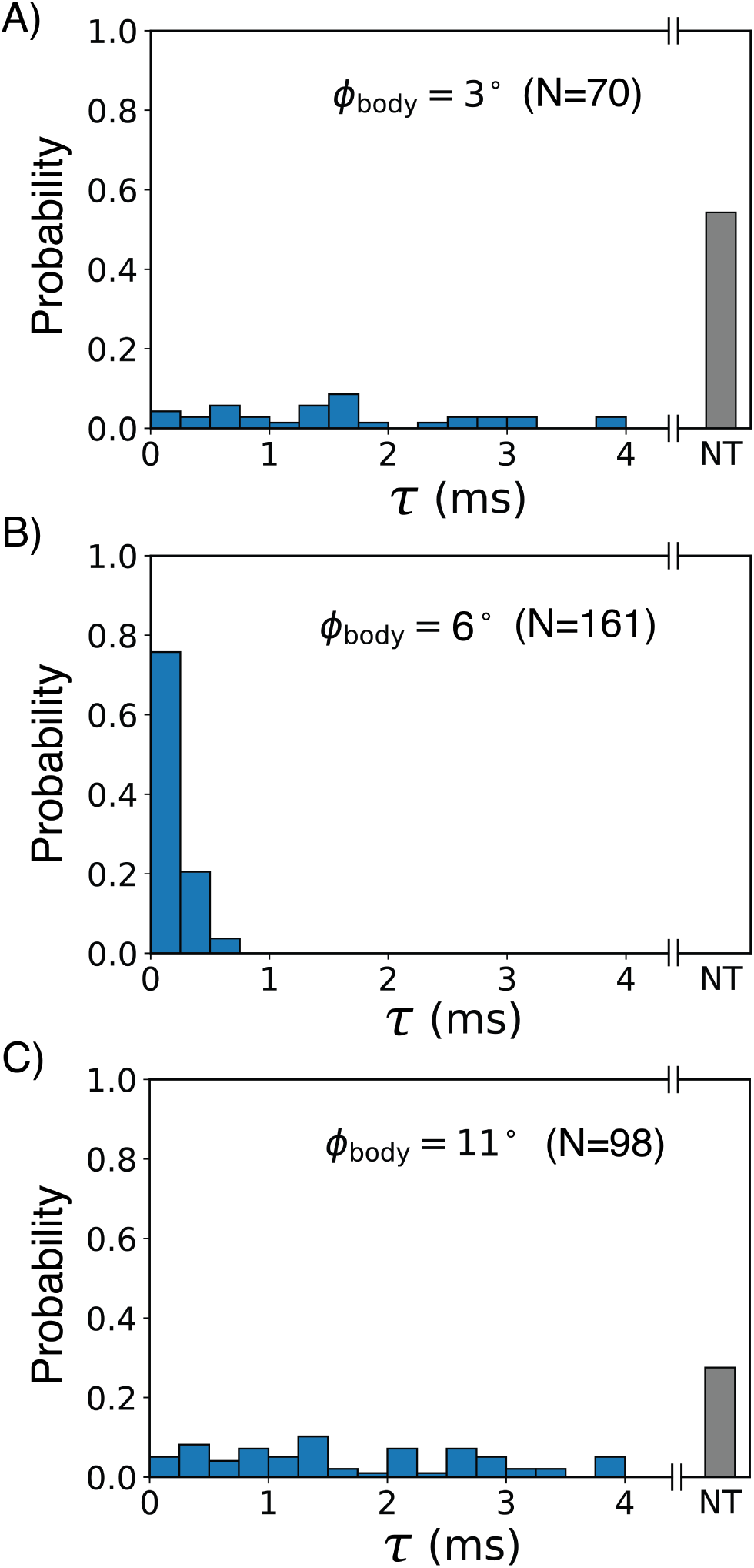
**Non-monotonic dependence of P/E tRNA hybrid formation on SSU body rotation**. The probability distribution of the time required to reach the P/E conformation, *τ*, shown for low, intermediate and high body rotation angles (*φ*_body_ = 3*^◦^*, 6*^◦^* and 11*^◦^*). N is the number of simulated runs for each rotation model. NT or “no transition” represents runs that did not reach the P/E state within the simulated timescale. A) The kinetics are slow for low rotation (*φ*_body_ = 3*^◦^*), where a majority of the runs did not reach the P/E state. B) The kinetics are faster at intermediate rotation (*φ*_body_ = 6*^◦^*), and almost all transitions occurred in less than one millisecond. C) At high body rotation (*φ*_body_ = 11*^◦^*), transitions occur over very long timescales, where a significant fraction did not reach the hybrid state.

The increase in P/E formation kinetics with initial body rotation may be understood in terms of accumulation and release of strain energy in the tRNA molecule. As the small subunit rotates, displacement of the anticodon stem loop (ASL; Fig. 4C) towards the E site leads to accumulation of internal energy in the tRNA. This increase in potential energy favors a transition to the P/E hybrid conformation, where displacement towards the E site allows the tRNA to relax. Overall, this strain accumulation and release process is similar to concepts used to describe conformational rearrangements in proteins^31^ and the idea that tRNA behaves as a (non-linear) “spring”.^32^ This spring-like interpretation can rationalize the observed increase in hybrid formation kinetics for body rotation angles up to ∼ 7*^◦^*. However, the strain mechanism would predict a monotonic relationship, which is incompatible with our results. Accordingly, non-monotonic behavior indicates that additional factors govern this relationship, such as non-trivial contributions of sterics and/or configurational entropy.

**Figure 4:**
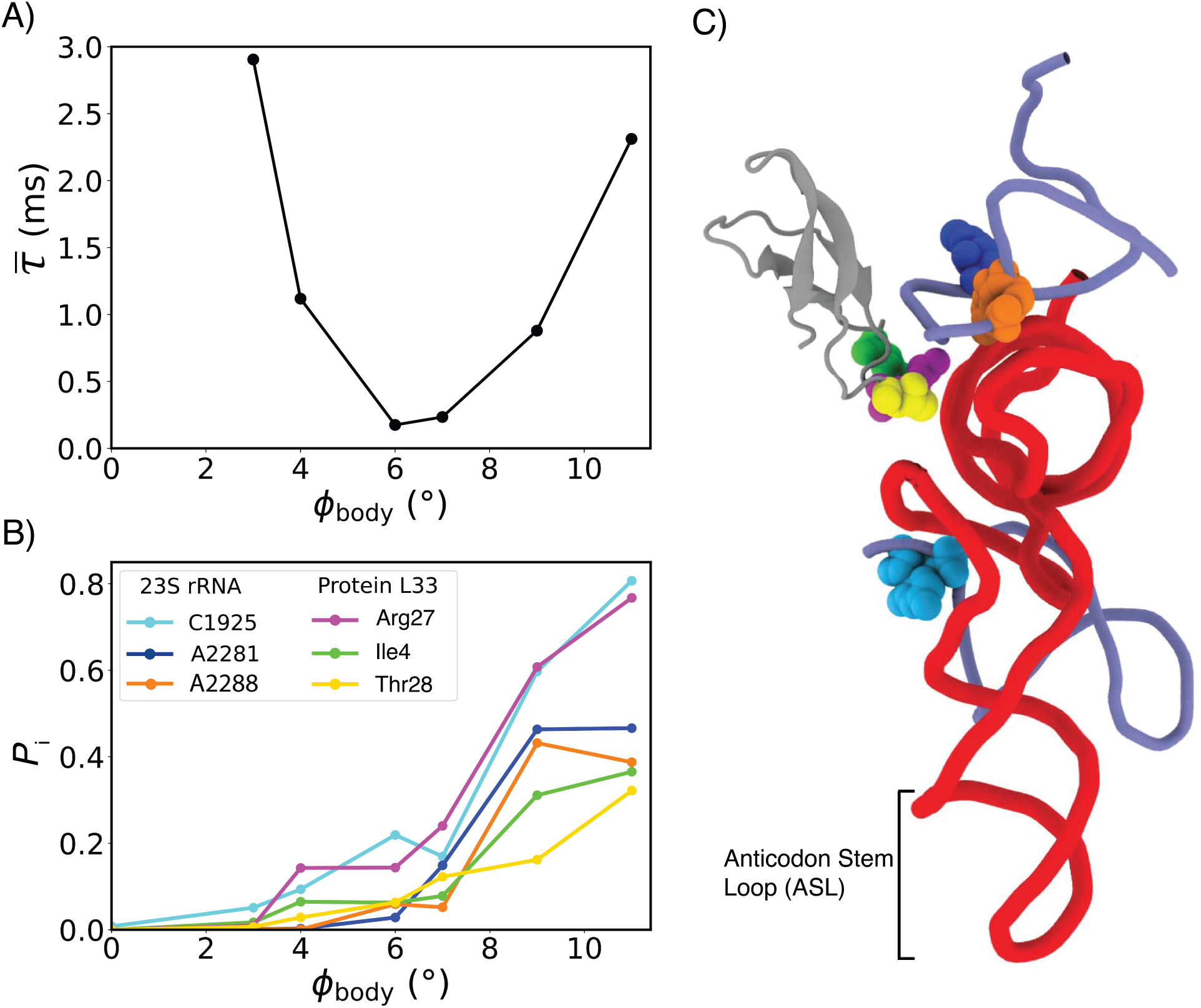
**Formation of steric interactions is correlated with slow down of tRNA kinetics**. A) There is a non-monotonic dependence of the mean first passage time *τ̄* for P/E hybrid formation on subunit rotation. The timescale for hybrid formation initially decreases with subunit rotation, with a minimum at *φ*_body_ = 6*^◦^*. For higher rotation angles, the timescale increases sharply. B) The probability that a ribosomal residue contacts P-site tRNA during a transition (*P*_i_), shown for six residues (i) that have increased contact probability at high rotation angles (L33 protein residues (Ile4, Arg27, Thr28); 23S rRNA residues C1925, A2281, A2288). There is a correlation between increased contact probabilities and the reduction in P/E formation kinetics. C) A simulated snapshot that highlights residues that form more steric interactions at higher rotation states.

### Rotation-dependent steric effects can rationalize tRNA kinetics

We next asked whether tRNA movement is controlled by rotation-dependent steric features in the ribosome. For this, we analyzed steric interactions that occur as the tRNA transitions between the P/P and P/E states. If formation of specific tRNA-ribosome interactions depends on the rotation state, then they may be responsible for hindering tRNA movement at higher rotation angles.

Through the analysis of tRNA-ribosome contacts, we identified specific ribosomal residues that have the potential to sterically hinder tRNA motion at higher body rotation angles. For this, we analyzed all simulated frames that were associated with a transition path (i.e. after release of the P site until binding of the E site. See SI for details.) and enumerated all ribosomal residues that formed short-range interactions with the tRNA when the SSU was fully rotated (11*^◦^*). 319 ribosomal residues were found to contact the tRNA in at least one simulated transition. We then calculated *P*_i_, the probability (fraction of simulated frames) that residue i was in contact with the tRNA while on a transition path. To identify candidate residues that could rationalize the predicted non-monotonic behavior, we applied the following two filters. First, *P*_i_ had to reach a relatively high value ( > 0.3). 17 residues satisfied this condition, including both rRNA and protein residues (see Figs. 4B and S3). Second, we considered residues for which *P*_i_ increased by at least a factor of two between 6*^◦^* and 11*^◦^*. This reduced the list of candidate residues to six (Figs. 4B and 4C), including three in the 23S rRNA (C1925, A2281 and A2288) and three in protein L33. The correlation between formation of steric interactions and rotation suggests that these specific residues may be responsible for impeding hybrid formation when the SSU is highly rotated.

### Protein L33 controls non-monotonic dependence of tRNA kinetics on subunit rotation

To directly test the influence of specific steric elements on tRNA dynamics, we introduced perturbations to our model and performed additional simulations of hybrid formation. For this, we considered three models in which protein L33 residues were modified/mutated and/or deleted. As described above, contact analysis in the original model (above) implicated protein L33 as a likely steric factor that can control tRNA dynamics at mid-to-high rotation angles. While some rRNA residues were also identified as relevant, we will focus our perturbation analysis on L33, since modifications in the model can mimic potential mutagenesis experiments. Accordingly, this analysis may be tested in future experimental studies.

We find that introducing modifications to the sterics of protein L33 almost fully eliminates the observed non-monotonic relationship between rotation and tRNA kinetics. For this discussion, we will refer to the original force field as Model O. To test the steric impacts of L33, we first considered a modified model in which two changes were introduced to L33 (Model Δ) : removal of the side chain of Arg27 and truncation of the N-terminal tail (Gly3, Ile4, Arg5; Fig. 5A). These changes are intended to represent an ideal experiment, where one could make a mutation (Arg to Gly) and truncation without introducing any indirect effects, such as changes in protein stability, binding or flexibility. We then used this model to perform additional simulations at varied degrees of rotation, from which the mean first passage time *τ̄* was calculated (see Table S3). We found that these modifications significantly altered the kinetics of hybrid formation at high body rotation angles (9 − 11*^◦^*). In contrast to the original model, the mean time for P/E formation exhibited almost no dependence on rotation (Figs. 5B). This is consistent with inferences based on contact analysis of Model O, and it confirms that the steric contributions of L33 are directly responsible for the observed non-monotonic dependence of P/E hybrid formation on subunit rotation.

**Figure 5:**
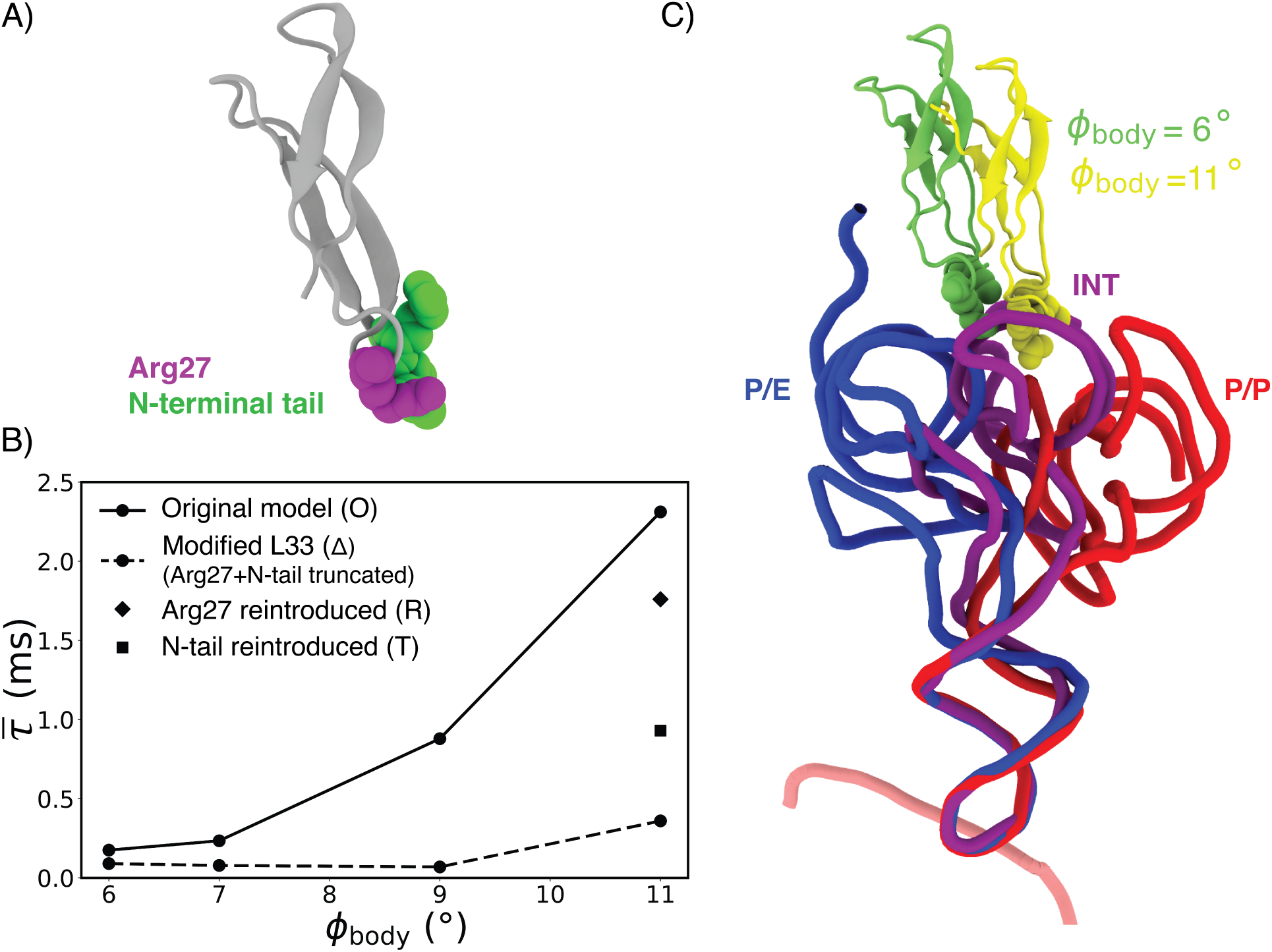
L33 protein acts as a molecular brake at high rotation angles. A) Structural depiction of protein L33 with Arg27 (magenta) and N-terminus tail (green) shown. B) Mean first-passage times for P/E formation *τ̄* as a function of SSU body rotation *φ*_body_. The unperturbed model (solid line, Model O) exhibits a non-monotonic dependence on rotation. Removing the side chain of Arg27 and the N-terminal tail (dashed line, Model Δ) eliminates this behavior. Simulations that reintroduced either Arg27 (Model R) or the N-terminal tail (Model T) reveal varying contributions to the kinetics. C) Structural snapshots of protein L33 when the ribosome is at *φ*_body_ = 6*^◦^* (green) and 11*^◦^* (yellow), which shows a rotation-dependent steric repositioning. Accordingly, rotation moves L33 inward, where it directly blocks the path of the tRNA. Multiple structures of tRNA illustrate the general movement that is associated with P/E formation (P/P, red; intermediate, purple; P/E, blue). Structures aligned based on the SSU.

To partition the steric contributions of Arg27 and the N-terminal tail, we considered two additional models in which the side chain of Arg27 (Model R) or the tail (Model T) was reintroduced. For each model, we performed independent simulations and calculated the corresponding mean first-passage time for the most highly-rotated state considered here (*φ*_body_ = 11*^◦^*). For ease of notation, we will refer to the mean time obtained with each model as *τ̄*_N_ (N=O, Δ, R, T). These calculations reveal complementary contributions to the underlying barrier (Figs. 5B). While Arg27 and the N-terminal tail both contribute to the apparent slowdown at high rotation, Arg27 has a larger contribution, where *τ̄*_R_ = 1.8ms and *τ̄*_T_ = 0.9ms. In terms of energetics, these two steric elements appear to make somewhat redundant contributions to the kinetics. To see this, we will assume there is a single dominant free-energy barrier, as has been obtained with a similar model applied to the T. thermophilus ribosome.^12^ We may then estimate the contribution of Arg27 to the free energy barrier as *k*_B_*T* ln(*τ̄*_R_*/τ̄*_Δ_) = 1.5*k*_B_*T*. Similarly, the contribution of the N-terminal tail would be given by *k*_B_*T* ln(*τ̄*_T_*/τ̄*_Δ_) = 0.8*k*_B_*T*. If these two contributions were independent and additive, then one would expect the barrier to be increased by 2.3*k*_B_*T* when the sterics of both are included. However, comparing the timescales when both are present or absent shows a smaller effect: *k*_B_*T* ln(*τ̄*_O_*/τ̄*_Δ_) = 1.7*k*_B_*T*. This indicates that these two elements serve as partially redundant steric attenuators that together lead to a pronounced non-monotonic relationship between tRNA movement and subunit rotation.

Structurally, Arg27 and the N-terminal tail forms a rotation-dependent steric barrier that increasingly obstructs tRNA movement as the body rotates. In the intermediate rotation states, these elements remain positioned away from the path of the tRNA. However, as body rotation increases beyond ∼ 7*^◦^*, the positioning of L33 places both Arg27 and the N-terminal tail directly in the path of the tRNA (Figs. 5C). At high rotation angles (9*^◦^*–11*^◦^*), this steric hindrance becomes pronounced, impeding tRNA progression and leading to the observed non-monotonic relationship between hybrid formation kinetics and body rotation. Removing these structural elements alleviates this barrier, explaining why their modification specifically eliminates the kinetic slowdown at high rotation angles. In terms of biological function, their redundant nature indicates that evolution has selected for RNA/protein sequences that ensure the dynamics of the ribosome are robust to minor changes.

## Discussion

This study reveals an intricate interplay between two major conformational motions in the ribosome. While previous studies have shown that subunit rotation contributes to hybrid formation, this relationship is complex, where the precise steric character of the ribosome modulates tRNA movement. This leads to the unexpected observation that rotation can increase (at mid-rotation) or decrease (at high rotation) the kinetics of tRNA movement. At a qualitative level, this non-monotonic behavior bears a resemblance to catch bonds, where bond lifetime increases with low applied force and then decreases at high force. In the ribosome, the rate of hybrid formation parallels a catch bond lifetime, where body rotation angle is responsible for introducing an applied force to the tRNA.

It is important to emphasize that the predicted contribution of rotation to tRNA dynamics should be robust to the precise energetics of the model. That is, while there is progress in understanding the ionic composition of the ribosome^33^ and its effects on conformational motions,^34^ the non-monotonic relationship originates from steric repulsion (i.e. excluded volume). Accordingly, this contribution should be present in more energetically-complete models, though the overall scale of the kinetics will certainly depend on the details of each model. Together, this suggests that efficient translation requires the ribosome to balance competing contributions of subunit rotation on tRNA dynamics.

## Methods

### Multi-basin all-atom structure-based model

Structure-based models describe effective energetics^35^ where experimentally known structures are defined to be potential energy minima. Here, we employed a multi-basin all-atom structure-based (SMOG) model^16^ to simulate P/E tRNA hybrid formation in the bacterial ribosome. For this, both pre-translocation (PRE) and post-translocation (POST) configurations are defined as potential energy minima (see SI Methods for definitions). We first generated single-basin SMOG models for pre- and post-translocation configurations of the ribosome. The interactions defined in these single basin models were then combined such that both the pre- and post-translocation configurations of the P-site tRNA were defined to be stable. In this composite model, every non-H atom is explicitly represented and is assigned unit mass. The potential energy is given by :

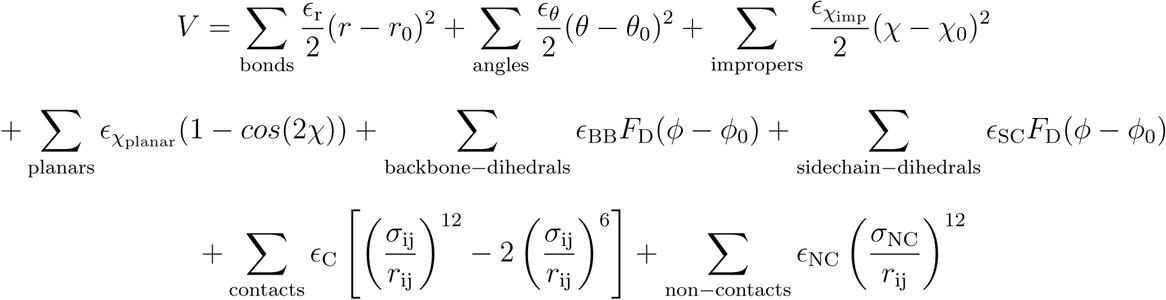

where

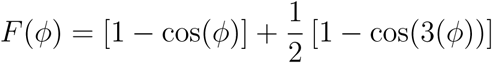

The model uses reduced energy units (*ϵ* = 1) with *ϵ* = 100*ϵ*/Å^2^, *ϵ* = 80*ϵ*/rad^2^, *ϵ* = 10*ϵ/*rad^2^ for improper dihedrals and *ϵ_χ_*= 40*ϵ*/rad^2^ for planar/ring dihedrals.

The covalent bond lengths (*r*_0_) and angles (*θ*_0_) are set based on the AMBER ff03 force-field.^36^ The contact lengths and dihedral angles parameters are defined such that a preassigned structure is defined as the potential energy minimum. Planar dihedral angles are described by a cosine function of periodicity 2 with minima at 0*^◦^* and 180*^◦^*. Consistent with previous implementations,^19^ the weight of each sidechain and backbone dihedral is set such that 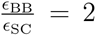. The Shadow Map algorithm^37^ with a cutoff of 6Å is used to identify the native contacts. Contact interactions are modeled as stabilizing 6-12 interactions. The native contact distance *σ*_ij_ is scaled by 0.96 to prevent the artificial expansion of the ribosome (measured by radius of gyration) due to thermal energy. Applying this scaling factor ensures that the radius of gyration of the simulated ribosome is consistent with the experimental structure.^17^ All atom pairs that are not in contact in the preassigned structure is given an excluded volume interaction with *σ*_NC_ = 2.5Å and *ϵ*_NC_ = 0.1*ϵ*. Contact energies are set such that 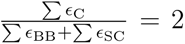 and Σ*ɛ*_C_ + Σ*ɛ*_BB_ + Σ*ɛ*_SC_ = Nc where N is the total number of atoms in the system. The force field is available on the smog-server force field repository (http://smog-server.org).

The model was constructed with all tRNA and mRNA contact and dihedral energies assigned based on the pre-translocation configuration. The multi-basin model was constructed such that the P-site tRNA has stabilizing contacts between both the P site and E site on the large subunit allowing the tRNA to undergo spontaneous hybrid formation. The stabilizing contacts between the P-site tRNA and E-site on the LSU were introduced from the posttranslocation structure leading the tRNA to have potential energy minima corresponding to both P/P and P/E states of the tRNA. Since SMOG model describes effective energetics,^35^ contacts that are formed transiently are assigned weaker strength (see SI Methods for details). The contacts formed by the tRNA with the LSU were assigned a weaker strength compared to the intra-ribosome contacts. Seven different SMOG models were generated, each corresponding to different SSU body rotation angles (*φ*_body_ = 0*^◦^*, 3*^◦^*, 4*^◦^*, 6*^◦^*, 7*^◦^*, 9*^◦^*, and 11*^◦^*). For each model, all intra-ribosome contacts and dihedrals were assigned based on a single rotated structure (see SI Methods for details).

### Simulation details

The structure-based force fields were generated using SMOG v2.4.^16^ All simulations were performed using OpenMM 8^38^ with the OpenSMOG^39^ libraries. All simulations were initiated from the pre-translocation A/A-P/P configuration. The simulations were performed at a constant temperature of 0.5 reduced units through use of Langevin dynamics protocols (drag term of 1/reduced time unit). For each representative rotation state, many independent simulations were performed (see Table S2). Simulations were terminated once the tRNA reached the E site of the LSU. Each simulation was continued for up to 2 billion timesteps, or 4×10^6^ reduced units. Yang et al.^40^ estimated that 1 reduced unit corresponds approximately to 1 ns, when simulating tRNA motion inside of the ribosome. To obtain this conversion factor, diffusion coefficients were calculated from simulations using a SMOG model and an explicit-solvent model.

### Structural metrics for P/E hybrid formation

We used the following structural coordinates to monitor the position of the tRNA molecule. *R*_E_: Distance between geometrical centers of the side chains of 23S rRNA residues G2421, C2422 (E site) and A76 of the P-site tRNA. *R*_E_ tracks the distance of the CCA tail of the P-site tRNA from the E site on the LSU. *R*_E_ is roughly 5nm and 0.2 nm in the P/P and P/E states.

*R*_FRET_: Distance between geometrical centers of the side chains of U47 in the A-site tRNA and U8 in the P-site tRNA. *R*_FRET_ is a commonly measured distance between labelled residues in the two tRNAs in sm-FRET experiments looking into tRNA dynamics. *R*_FRET_ reports on tRNA elbow movement, and it is ∼ 4.3nm and ∼ 6.3nm in the P/P and P/E states.

### Ribosome Angle Decomposition (RAD) coordinates

The SSU body and head undergo various modes of rotational motions during the elongation cycle. The Ribosome Angle Decomposition (RAD) method^41^ provides a complete coordinate system to describe the orientation of ribosomal small subunit body and head. Any orientation of the SSU body is reported in terms of rotation angle *φ*_body_ about a fixed axis 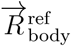, tilt-like rotation *θ*_body_ about the calculated axis *T⃗*_body_, the tilt direction *ψ*_body_ and a translation vector Δ*x⃗*_body_. Here the orientation of the SSU body is described with respect to the LSU. The orientation of the head domain is described using analogous coordinates (*φ*_head_, *θ*_head_, *ψ*_head_ & Δ*x⃗*_head_). To decompose the orientations of body and head in terms of RAD coordinates, first the core residues in the ribosomal large subunit, small subunit body, and head are identified through structure alignment steps with respect to reference E. coli structures. To identify the core residues, STAMP^42^ is used for initial structure alignment, followed by a pruning algorithm which gets the root mean square deviation of the core with respect to reference E. coli structure to be ∼ 1Å. Once the core has been identified, rigid body approximation is applied to calculate the Euler angles for both SSU body and head. While calculating rotation angles for the simulated trajectories, to ensure consistent analysis across all seven rotation states, we identified the common core residues that were present in all seven representative structures.

### Selection of representative rotation structures

RAD was utilized to identify representative rotation structures that span the full range of body rotation angles. To focus specifically on the impact of body rotation on hybrid formation, we selected structures with small contributions from other modes of ribosome dynamics. We considered the rotational orientations of all published E. coli ribosome structures from the RCSB database (resolution of 8Å or greater; 412 structures). From this, we identified representative structures ranging from unrotated (∼ 0*^◦^*) to highly-rotated (∼ 11*^◦^*) states, where all other rotational modes are small. Specifically head rotation, head tilt and body tilt were set to less than 4*^◦^*. The following reference structures were used (RCSB accession codes given): 0.0*^◦^*(4V9D^43^), 3.2*^◦^*(6WDI^44^), 4.6*^◦^*(6GXN^45^), 6.0*^◦^*(4V6E^46^), 7.8*^◦^*(6GXP^45^), 9.3*^◦^*(4V9D^43^), and 10.4*^◦^*(4V7D^47^).

### Original and modified L33 models

To investigate the steric contribution of specific ribosomal elements on tRNA dynamics, we considered multiple structural models of the ribosome. These models were constructed by systematically modifying protein L33, particularly its Arg27 residue and the N-terminal tail (Gly3, Ile4, and Arg5).

- Model O (Original Model): The unperturbed ribosome model, where all native contacts and dihedrals for L33 were retained. This model includes full steric contributions from Arg27 and the N-terminal tail of L33. Simulations were continued for up to 4ms, except for *φ*_body_ = 0*^◦^* (8ms).
- Model Δ (Modified Model): In this model, steric perturbations were introduced to protein L33. The side chain of Arg27 was removed, and the N-terminal tail residues Gly3, Ile4, and Arg5 were truncated. Simulations were continued for up to 4ms.
- Model R (Arg27 reintroduced): To isolate the contribution of Arg27, this model reintroduced (relative to Model Δ) the side chain of Arg27, while the N-terminal tail was absent. This enables evaluation of the specific steric effects of Arg27 on tRNA movement. Maximum timescale simulated with this model is 2.4ms.
- Model T (N-terminal tail reintroduced): In this model, the N-terminal tail of L33 (Gly3, Ile4, and Arg5) was reintroduced, while the side chain of Arg27 was absent. Maximum timescale simulated with this model is 2.4ms.

## Acknowledgement

This work was supported by the National Institutes of Health (grant R35GM153502-01) and the National Science Foundation (grant MCB-1915843). Work at the Center for Theoretical Biological Physics was supported by the National Science Foundation (grant PHY-2019745). We thank AMD for the donation of critical hardware and support resources from its HPC Fund that made this work possible.

