## Supplementary Information for "How subunit rotation controls tRNA dynamics in the ribosome"

### Supplementary Information for:
### How subunit rotation controls tRNA dynamics in the ribosome

Sandra Byju[1,2] and Paul C. Whitford[1,2]

[1] *Center for Theoretical Biological Physics, Northeastern University, 360 Huntington Ave, Boston, MA 02115, USA*
[2] *Department of Physics, Northeastern University, Dana Research Center 111, 360 Huntington Ave, Boston, MA 02115, USA*

#### List of Tables

S1 Structure and simulation details of representative rotation states . . . . . S2
S2 Simulation results for P/E hybrid formation with different rotation states (Model O). . . . . S3
S3 Simulation results for P/E hybrid formation at different rotation states with Model Δ. . . . . S3

#### List of Figures

S1 POST translocation structure . . . . . S4
S2 Distribution of transition time $\tau$ for $\phi_{\text{body}} = 4°$, $7°$ and $9°$ . . . . . S5
S3 Interaction probabilities of identified ribosomal residues with tRNA during P/P to P/E transition across body rotation angles . . . . . S6

#### Contents

**1 SI Methods** **S7**
1.1 Preparation of PRE and POST translocation configuration . . . . . S7
1.1.1 A-site Finger . . . . . S7
1.1.2 Residue 1847 in 23S rRNA . . . . . S7
1.1.3 L1 protein . . . . . S7
1.1.4 L10 protein . . . . . S8
1.1.5 P-site tRNA, A-site tRNA and mRNA in an A/A-P/P configuration (PRE translocation) S8
1.1.6 P-site tRNA, E-site tRNA and mRNA to define a P/P-E/E configuration (POST translocation) . . . . . S8
1.2 Multi-basin model parameters . . . . . S9
1.3 Generation of representative rotated models . . . . . S9
1.4 Calculation of P/P to P/E transition time . . . . . S9
1.5 Additional rotation models . . . . . S10
1.6 Analysis of tRNA-ribosome steric interactions during transition path . . . . . S10

| Experimental Structures Used | | | | | | | Simulation Details | |
|---|---|---|---|---|---|---|---|---|
| PDB ID | $\phi_{\text{body}}(^\circ)$ | $\theta_{\text{body}}(^\circ)$ | $\phi_{\text{head}}(^\circ)$ | $\theta_{\text{head}}(^\circ)$ | Xray/EM | Res(Å) | $\langle\phi_{\text{body}}\rangle(^\circ)$ | label |
| 4V9D | 0.0 | 0.0 | 0.0 | 0.0 | Xray | 3.0 | 0.0 | 0 |
| 6WDI | 3.2 | 0.8 | -1.0 | 1.6 | EM | 4.0 | 3.0 | 3 |
| 6GXN | 4.6 | 2.1 | 1.0 | 2.1 | EM | 3.9 | 3.9 | 4 |
| 4V6E | 6.0 | 1.2 | -0.4 | 1.5 | Xray | 3.7 | 5.5 | 6 |
| 6GXP | 7.8 | 1.7 | 3.8 | 3.8 | EM | 4.4 | 7.2 | 7 |
| 4V9D | 9.3 | 0.0 | 2.7 | 1.7 | Xray | 3.0 | 9.2 | 9 |
| 4V7D | 10.4 | 2.6 | 3.3 | 3.0 | EM | 7.6 | 10.6 | 11 |
| 4V7C | 9.5 | 1.9 | 2.7 | 2.2 | EM | 7.6 | 9.4 | 9 |
| 7PJT | 9.9 | 2.1 | 3.2 | 2.5 | EM | 6.0 | 10.3 | 10 |

Table S1: Summary of structure and simulation details of representative rotation states. Columns include PDB IDs of the reference structures, body rotation ($\phi_{\text{body}}$), body tilt ($\theta_{\text{body}}$), head rotation ($\phi_{\text{head}}$), head tilt ($\theta_{\text{head}}$), structural determination method (X-ray crystallography or cryo-EM), resolution of the structure (Res). $\langle\phi_{\text{body}}\rangle$ average rotation angle when the given structure was used to define the intra-ribosome interactions in the SMOG model. The final column lists the label used to refer to set of simulations with the given rotation state modeled, and it is the average simulated rotation angle rounded to the nearest integer. All subsequent references to rotation angles use these rounded values. For completeness, two additional rotation models (4V7C and 7PJT) were used to perform simulations.

| PDB ID | $\phi_{\mathrm{body}}(^\circ)$ | Total runs N | $\mathrm{N}_{\mathrm{transition}}$ | $\bar{\tau}$ (ms) |
|---|---|---|---|---|
| 4V9D | 0 | 4 | 0 | 8.00 |
| 6WDI | 3 | 70 | 32 | 2.90 |
| 6GXN | 4 | 119 | 118 | 1.12 |
| 4V6E | 6 | 161 | 161 | 0.18 |
| 6GXP | 7 | 132 | 132 | 0.23 |
| 4V9D | 9 | 123 | 119 | 0.88 |
| 4V7D | 11 | 98 | 75 | 2.31 |
| 4V7C | 9 | 109 | 98 | 1.73 |
| 7PJT | 10 | 108 | 105 | 0.98 |

Table S2: Summary of simulation results for P/E hybrid formation at different body rotation states, performed with the original model (Model O). Columns include the PDB ID of the reference structure, the rotation state $\phi_{\mathrm{body}}$, the total number of independent simulations N, the number of runs in which the tRNA molecule reached the P/E state within the simulated timescale $\mathrm{N}_{\mathrm{transition}}$, and the mean first passage time $\bar{\tau}$ for P/E hybrid formation. All simulations were performed for a maximum time of 4ms, except for $\phi_{\mathrm{body}} = 0^\circ$ (8ms).

| PDB ID | $\phi_{\mathrm{body}}(^\circ)$ | Total runs N | $\mathrm{N}_{\mathrm{transition}}$ | $\bar{\tau}$ (ms) |
|---|---|---|---|---|
| 4V6E | 6 | 184 | 184 | 0.09 |
| 6GXP | 7 | 114 | 114 | 0.08 |
| 4V9D | 9 | 107 | 107 | 0.07 |
| 4V7D | 11 | 112 | 112 | 0.36 |
| 4V7C | 9 | 146 | 146 | 0.25 |
| 7PJT | 10 | 142 | 142 | 0.21 |

Table S3: Simulation results for P/E hybrid formation kinetics using the modified Model $\Delta$, in which protein L33 was altered by removing the side chain of Arg27 and truncating the N-terminal tail (Gly3, Ile4, Arg5). Columns include the PDB ID of the reference structure, the rotation state $\phi_{\mathrm{body}}$, the total number of independent simulations N, the number of runs in which tRNA transitioned to the P/E state within the simulated timescale $\mathrm{N}_{\mathrm{transition}}$, and the mean first passage time $\bar{\tau}$ for P/E hybrid formation.

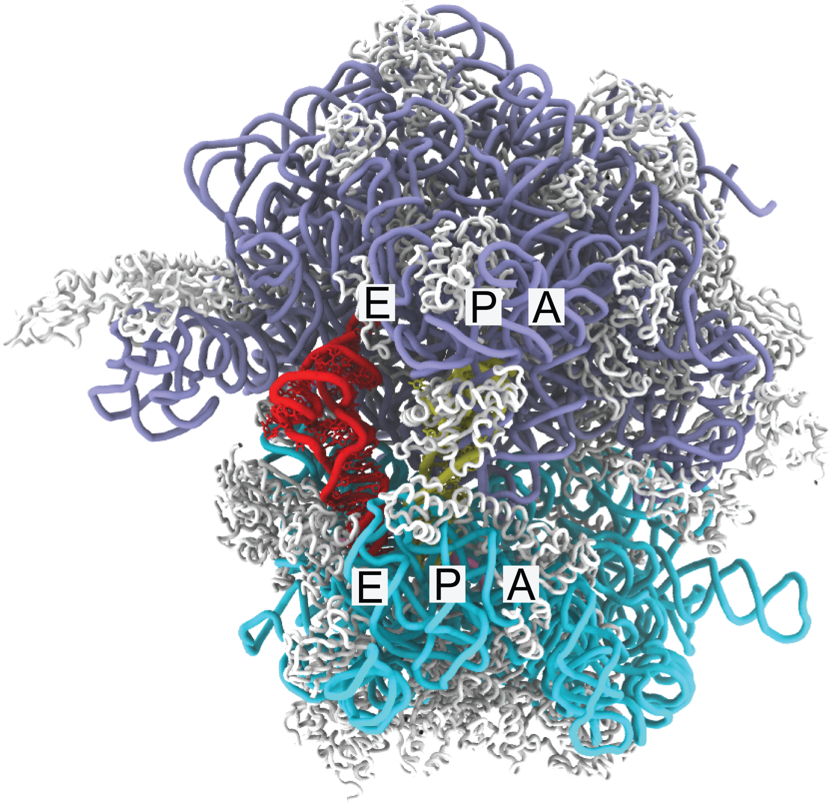


Figure S1: **POST translocation structure** E. coli ribosome (PDB ID: 4V9D[1]) in the POST-translocation state with the ribosome in the classical unrotated configuration. The P-site (yellow) and E-site (red) tRNAs are in the P/P and E/E conformations, respectively. The LSU rRNA (23S and 5S) is shown in purple, SSU rRNA (16S) in cyan, proteins in white and mRNA in pink.

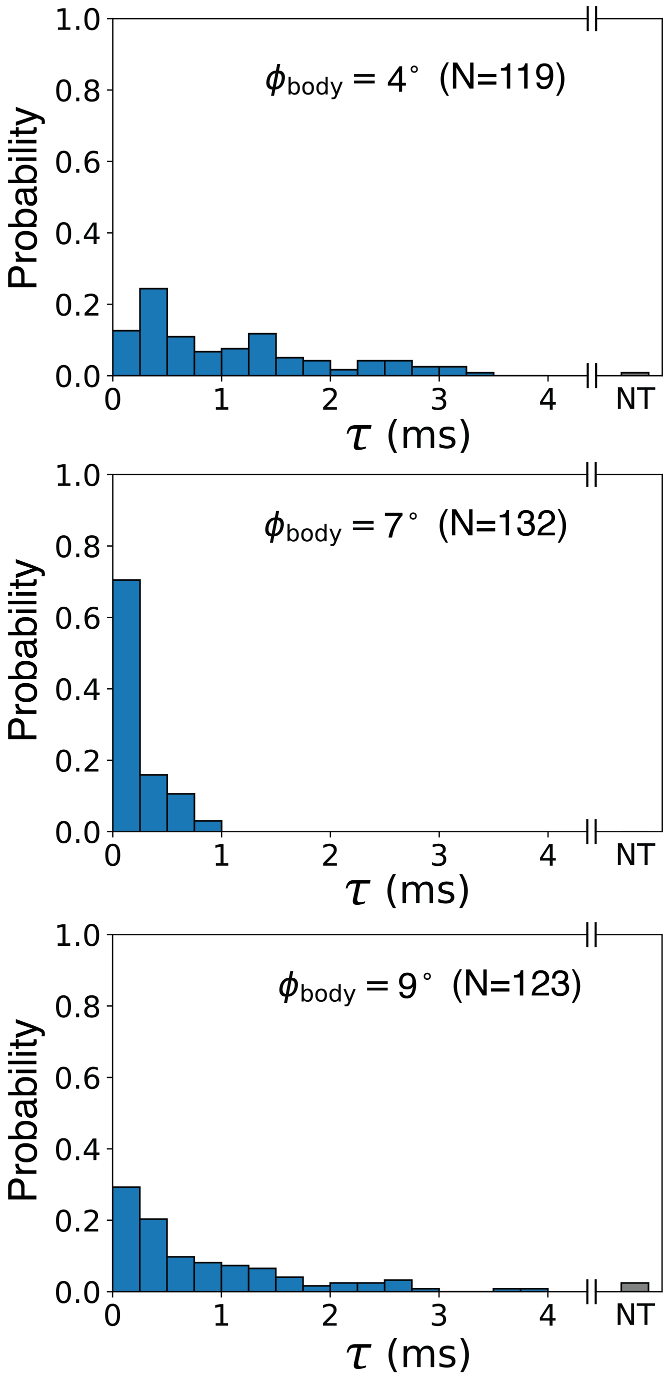


Figure S2: **P/E formation timescales** The probability of P/P to P/E transition times $\tau$ is shown for rotation states $\phi_{\text{body}} = 4°$, $7°$ and $9°$. N is the total number of simulated runs. NT or "no transition" represent fraction of runs which did not reach the P/E state within the simulated timescale (simulation capped at 4 ms).

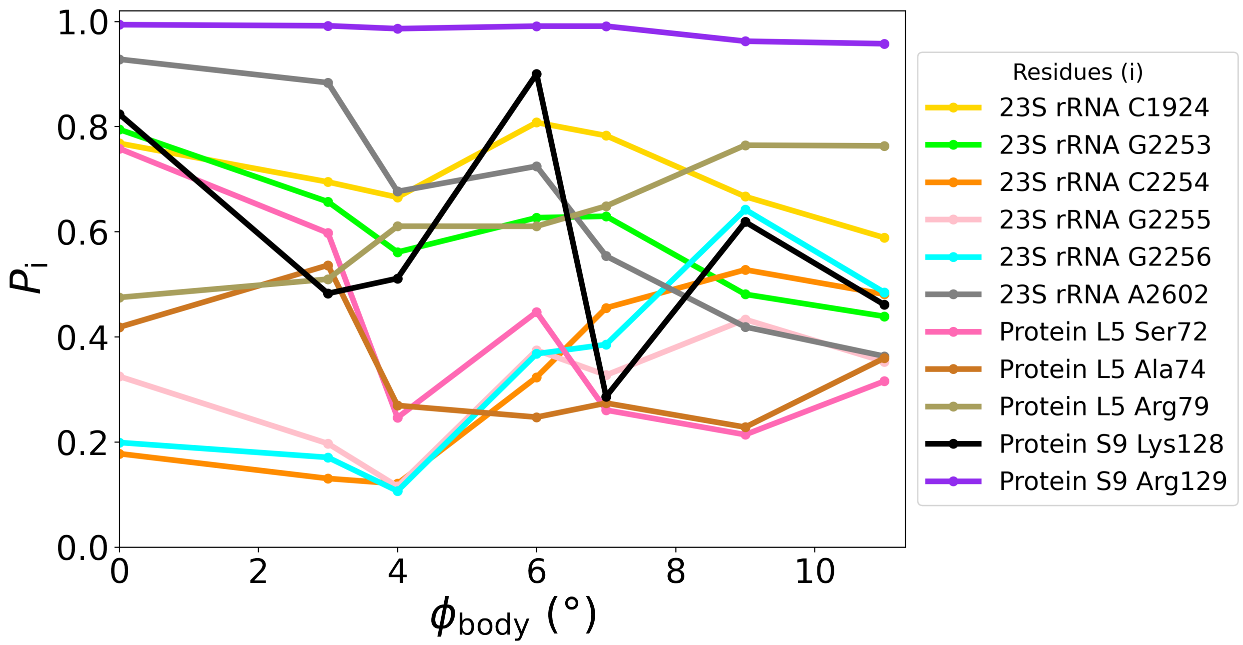


Figure S3: **Interaction probabilities of ribosomal residues with tRNA during P/P to P/E transition path across body rotation angles.** Interaction patterns for the remaining 11 residues (i) of the 17 residues that show high probabilities of interacting with tRNA at 11° body rotation but lack a substantial increase relative to 6°.

### 1 SI Methods

#### 1.1 Preparation of PRE and POST translocation configuration

To define the PRE and POST translocation states, we used PDB ID: 4V9D structure[1], which provides a high-resolution snapshot (3Å) of the unrotated bacterial ribosome with a tRNA in the classical P/P state. However, some structural elements are missing in this structure, which necessitated additional structural modeling. Specifically, this structure does not include the A-site and E-site tRNAs, Helix 38 (A-site finger), residue 1847 in 23S rRNA, or proteins L1 and L10. To address these limitations, structural information was incorporated from other high-resolution bacterial ribosome structures that contain the missing components. All the ions resolved in the structure including $Mg^{2+}$ and $Zn^{2+}$ were removed.

##### 1.1.1 A-site Finger

To incorporate the missing A-site finger residues 886 to 891 in the 23S rRNA, structural information was adopted from the unrotated ribosome PDB ID: 7K00[2], which provides the coordinates for this region. The following protocol was followed:

1. **Alignment of coordinates:** The 23S rRNA coordinates from 7K00, which include residues 886 to 891, were aligned to the 4V9D structure, where alignment was performed based on residues immediately upstream and downstream of the missing region (883, 884, 885, 892, 893, and 894).

2. **Translation and harmonic restraints application to resolve clashes:** The aligned missing region (residues 886 to 891) was manually translated several angstroms away from its initial position to prevent atomic clashes with the existing A-site finger structure and surrounding residues. Harmonic restraints were then applied to every atom in the translated missing region, with minima set to the coordinates of the aligned structure from 7K00.

3. **Energy minimization:** Energy minimization was conducted using a structure-based potential (SMOG). During this process:

   - All atoms outside the missing region were fixed to their original positions.
   - The excluded volume term of the potential prevented the introduction of atomic clashes during minimization. Energy minimization allowed the missing region to relax into the aligned configuration, ensuring proper incorporation into the model while maintaining steric compatibility.
   - Energy minimization was sequentially performed with harmonic restraint strengths of 10, 50, and 100 $\epsilon/\text{nm}^2$.
   - Finally, the system was energy minimized without position restraints while fixing all atoms except for the A-site finger.

##### 1.1.2 Residue 1847 in 23S rRNA

The procedure for incorporating the missing residue 1847 in the 23S rRNA followed the same steps as outlined for residues 886 to 891 in the A-site finger. Structural information for residue 1847 was adopted from PDB ID: 7K00[2]. The residues immediately upstream and downstream of the missing region used for alignment include residues 1845, 1846, 1848 and 1849.

##### 1.1.3 L1 protein

The procedure for incorporating the missing protein L1 in the LSU followed the same general steps as outlined for residues 886 to 891 in the A-site finger. Structural information for protein L1 was adopted from PDB ID: 4V7C[3]. The residues 2095 to 2192 of the 23S rRNA were used as reference for alignment.

##### 1.1.4 L10 protein

The procedure for incorporating the missing protein L10 in the LSU followed the same general steps as outlined for residues 886 to 891 in the A-site finger. Structural information for protein L10 was adopted from PDB ID: 4V7C[3]. The residues 1052 to 1102 and 1107 to 1116 of the 23S rRNA were used as reference for alignment.

##### 1.1.5 P-site tRNA, A-site tRNA and mRNA in an A/A-P/P configuration (PRE translocation)

To incorporate the P-site tRNA, A-site tRNA, and mRNA, structural information was remodeled using coordinates from PDB ID: 4V6F[4]. The 4V6F structure of the unrotated ribosome contains all three tRNAs (A, P and E) with a single modified residue (MIA37) and polyU mRNA. The following steps were specific to this incorporation:

1. A-site tRNA: A-site tRNA was not resolved on the 4V9D structure. The alignment of the tRNA was performed using residues within 3Å of the A-site tRNA in 4V6F as reference. These residues are conserved and present in 4V9D.

2. P-site tRNA: The P-site tRNA in 4V9D (excluding residue 37) was used as a reference for alignment. The P-site tRNA from 4V6F was then aligned to this reference. The P-site tRNA in 4V9D was removed to ensure consistency. The 4V6F structure contains all three tRNAs (A, P, and E) with a uniform structure, including the modified residue MIA37.

3. mRNA: Alignment was similarly performed using residues within 3Å of the mRNA in 4V6F. Residues 42 to 55 from mRNA of 4V6F were retained after alignment. The mRNA already present in 4V9D was removed.

4. No conflicting contacts with the neighboring atoms were observed after alignment. The system was energy minimized with all other components fixed except for A-site tRNA, P-site tRNA and mRNA based on SMOG structure-based potential, and no harmonic restraints were applied.

5. All missing terminal oxygen atoms of proteins and OP3 atoms at the 5' terminal end of rRNA chains were added manually.

##### 1.1.6 P-site tRNA, E-site tRNA and mRNA to define a P/P-E/E configuration (POST translocation)

To incorporate the P-site tRNA, E-site tRNA and mRNA, structural information was remodeled using coordinates from PDB ID: 4V6F[4]. The 4V6F structure of the unrotated ribosome contains all three tRNAs (A, P and E) with a single modified residue (MIA37) and polyU mRNA. The following steps were specific to this incorporation:

1. P-site tRNA: The P-site tRNA in 4V9D (excluding residue 37) was used as a reference for alignment. The P-site tRNA from 4V6F was then aligned to this reference. The P-site tRNA in 4V9D was removed to ensure consistency. The 4V6F structure contains all three tRNAs (A, P, and E) with a uniform structure, including the modified residue MIA37.

2. E-site tRNA: The E-tRNA was aligned by matching residues within 3Å of the E-site tRNA in 4V6F, which are conserved and also present in 4V9D. During alignment, two conflicting contacts were identified between the E-site tRNA and the S7 protein. To resolve these conflicts, energy minimization was performed with all atoms fixed except for the S7 protein and all E-site tRNA–S7 contacts removed from the SMOG structure-based potential.

3. mRNA: Alignment was similarly performed using residues within 3Å of the mRNA in 4V6F. Residues 39 to 52 from mRNA of 4V6F were retained after alignment. The mRNA already present in 4V9D was removed.

4. The system was energy minimized with all other components fixed expect P-site tRNA, E-site tRNA and mRNA based on SMOG structure-based potential, and no harmonic restraints were applied.

5. All missing terminal oxygen atoms of proteins and OP3 atoms at the 5' terminal end of rRNA chains were added manually.

#### 1.2 Multi-basin model parameters

The contacts formed by the tRNA with the LSU were assigned a weaker strength compared to the intra-ribosome contacts. The contacts between the P-site tRNA with the P site were assigned a contact strength of $0.05\epsilon_{\mathrm{C}}$. The contact strength for contacts between tRNA and E site from the post-translocation configuration were set to $0.25\epsilon_{\mathrm{C}}$. To ensure stable binding of the tRNA with the E site during hybrid formation, the contacts between A76 of CCA tail and residues G2421/C2422 of the 23S rRNA (LSU) were assigned a stronger contact strength of $1.5\epsilon_{\mathrm{C}}$. To prevent the unfolding of the the tRNA which was observed in the pre-production runs, harmonic restraints with spring constant of $200\epsilon/\mathrm{nm}^2$ were introduced between atom pairs of residues G1 and C72 that formed a contact. This prevented the unfolding of the tRNA acceptor region arising due to low contact density in this region when using this model.

#### 1.3 Generation of representative rotated models

Seven different SMOG models were generated, each corresponding to rotation of $\phi_{\mathrm{body}} = 0°$, $3°$, $4°$, $6°$, $7°$, $9°$, and $11°$. For each model, all intra-ribosome contacts and dihedrals were assigned based on a representative rotated structure. To overcome the challenge of missing regions in experimentally resolved structures and to ensure consistent atomic representation across all models, we had to apply a systematic modeling strategy. Starting with the classical unrotated ribosome structure (PDB ID: 4V9D), we generated each rotated structure by applying position restraints on common backbone rRNA atoms and all common protein atoms using the corresponding representative rotation structure (Table S1) as reference. The refinement protocol began with energy minimization using a position restraint strength of $k = 1\ \epsilon/\mathrm{nm}^2$. While maintaining these position restraints, the system was then subjected to simulated annealing where the temperature was gradually decreased from $T = 0.25$ to 0 reduced units, allowing local structural elements to relax while preserving the target rotated state. This refinement process was then repeated with stronger position restraint strengths of $k = 10$ and $100\ \epsilon/\mathrm{nm}^2$. For cases where entire peripheral ribosomal proteins (e.g., L1 and L10) were missing in the reference structure, these elements were remodeled after the refinement process as described earlier. This approach ensured that all seven models maintained identical atomic composition, with differences solely attributed to the degree of body rotation.

#### 1.4 Calculation of P/P to P/E transition time

We calculated the transition time $\tau$ for the movement of the tRNA from the P/P state to the P/E hybrid state as the time between when the tRNA is released from the P site and when it reached the P/E state using structural coordinates $R_{\mathrm{E}}$ and $R_{\mathrm{FRET}}$.

1. All simulations were initiated from the PRE-translocation state, where the tRNA was in the classical P/P conformation.

2. Release of tRNA from the P site was determined by monitoring the distance between residue C75 in the 3'-CCA tail and the ribosomal P site (23S rRNA residue 2251). A threshold distance of 6 Å was applied to classify when the tRNA has released the P site.

3. The time of arrival at the E site was determined by calculating $R_{\mathrm{E}}$ (the distance between residue A76 of the tRNA and the E site residues 2421, 2422). Applied a threshold of 0.2 nm to indicate that the tRNA reached the E site. Additionally, we calculated $R_{\mathrm{FRET}}$, the distance between the elbow region of the P-site tRNA and the A-site tRNA. The tRNA was defined as having reached the E site when $R_{\mathrm{E}} < 0.2$ nm and $R_{\mathrm{FRET}} > 6.3$ nm.

4. Determined the mean first passage time $\bar{\tau}$ for each rotation angle ($\phi_{\text{body}}$) as $\bar{\tau} = \frac{\sum_{i=1}^{N} \tau_i}{N}$, where $N$ is the total number of simulated runs.

5. Simulations where the tRNA did not reach the P/E state were included in time averages by using the maximum simulated time for $\tau_i$.

#### 1.5 Additional rotation models

To confirm that the dynamics for high rotation angles did not depend on the specific structural model that was used, we performed simulations with two additional rotation models $\phi_{\text{body}} = 9°$ (PDB ID: 4V7C [3]) and $\phi_{\text{body}} = 10°$ (PDB ID: 7PJT [5]). Models O and $\Delta$ for these rotation states were constructed using the same methodology described previously. For the $\phi_{\text{body}} = 9°$ Model O (4V7C), 98 out of 109 simulations reached the P/E state, with a mean first passage time ($\bar{\tau}$) of 1.73 ms. Similarly, for the $\phi_{\text{body}} = 10°$ Model O (7PJT), 105 out of 108 simulations transitioned to the P/E state, with $\bar{\tau}$= 0.98 ms (Table S2). Both models exhibited slower kinetics compared to intermediate rotation angles ($\phi_{\text{body}} = 6°$ and $\phi_{\text{body}} = 7°$), further supporting the non-monotonic relationship between SSU body rotation and P/E hybrid formation kinetics. For the L33-modified Model $\Delta$, $\bar{\tau}$ for P/E formation was significantly reduced to 0.25 ms ($\phi_{\text{body}} = 9°$) and 0.21 ms ($\phi_{\text{body}} = 10°$) (Table S3). These results reinforce the conclusion that modifying L33 by removing steric elements effectively eliminates the non-monotonic dependence of hybrid formation kinetics on SSU rotation. Including these additional rotation models complements and validates the observed trends, further underscoring the role of L33 sterics in controlling tRNA dynamics at high rotation angles.

#### 1.6 Analysis of tRNA-ribosome steric interactions during transition path

To analyze the steric interactions between the tRNA and ribosome during transitions from the P/P to P/E states at high rotation angle, we identified all ribosomal residues that formed short-range interactions with the tRNA while on a transition path. As described in Sec. 1.4, a transition path was defined as the portion of the trajectory between the release of the tRNA from P site and its subsequent binding to the E site. During a transition event, all ribosomal residues that formed short-range contacts with the tRNA were identified. A contact was defined as any residue within 4 Å of the tRNA. For each residue, the probability of contact ($P_{\text{i}}$) was calculated as the fraction of frames within the transition paths where the residue was within 4 Å of the tRNA.
